# Novel application method for mesenchymal stem cell therapy utilizing its attractant–responsive accumulation property

**DOI:** 10.1101/626275

**Authors:** Nobuyuki Ueda, Ikiru Atsuta, Yasunori Ayukawa, Takayoshi Yamaza, Akihiro Furuhashi, Ikue Narimatsu, Yuri Matsuura, Ryosuke Kondo, Yu Watanabe, Xiaoxu Zhang, Kiyoshi Koyano

**Affiliations:** Section of Implant and Rehabilitative Dentistry, Division of Oral Rehabilitation, Faculty of Dental Science, Kyushu University, Fukuoka, Japan; Division of Advanced Dental Devices and Therapeutics, Faculty of Dental Science, Kyushu University, Fukuoka, Japan; Department of Molecular Cell and Oral Anatomy, Faculty of Dental Science, Kyushu University, Fukuoka, Japan

**Keywords:** Mesenchymal stem cell, tooth extraction socket, collagen gel, systemic administration, wound healing, scaffold

## Abstract

**Objectives:** It remains difficult to control the delivery of appropriate amounts of mesenchymal stem cell (MSC)-based cell therapies. To examine the ability of MSCs to accumulate at sites of damage and potential therapeutic benefit of providing continuous migration of MSCs to these sites, we observed the effect of MSCs administered in a collagen gel scaffold on healing of a tooth extraction site.

**Materials and Methods:** MSCs isolated from the bone marrow of green fluorescent protein (GFP)-expressing donor mice were expanded for 3 weeks in three-dimensional (3-D) culture using a collagen gel scaffold, and evaluated to confirm the efficacy of the scaffold. Next, MSCs suspended in collagen gel were subcutaneously administered into the backs of mice. Two days later, extraction of the maxillary first molar was carried out. Numbers of MSCs in scaffolds, migration and accumulation around the extracted tooth cavity, extraction site healing, and presence of MSCs in vital organs were evaluated.

**Results:** MSCs cultured in the collagen gel scaffold maintained stemness for 2 weeks. After subcutaneous administration, numbers of MSCs in scaffolds slightly decreased over time, but cells survived for at least 2 weeks. After tooth extraction, GFP-expressing MSCs were confirmed in the surrounding mucosa of the extracted tooth cavity; in the scaffold group, numbers of MSCs increased over time and fewer were observed in lung tissue. Wound healing was enhanced by injection of MSCs via the tail vein or into the back compared with the untreated control group.

**Conclusions:** Delivery in a collagen gel could maintain the characteristics of MSCs, which migrated to the damaged area and promoted wound healing without side effects occurring with conventional administration methods.

## Introduction

Mesenchymal stem cells (MSCs), which are known to contribute to tissue regeneration and repair [1], are normally present in a dormant state in almost all organs [2]. Upon the release of cytokines by a stimulus such as inflammation or tissue injury, MSCs begin to proliferate and migrate to the site from where cytokines are being released; i.e. the damaged area [3]. However, proliferation and migration of MSCs takes time, and a lack of MSCs at required sites may impede tissue healing. Although some methods utilize the regenerative properties of stem cells for therapy [4], in terms of cell delivery methods, a gold standard has not been established and further study is needed. Upon systemic delivery of stem cells through blood circulation, some become clogged in the lung, where they can cause pulmonary embolism [5]. For topical applications, controlling and stabilizing stem cells in the desired area is not always accomplished, and if cells are too dense, they undergo apoptosis [6].

Therefore, we focused on the potential of MSCs to autonomously accumulate in damaged tissue. Our hypothesis is that subcutaneous administration of a mixture of scaffold and MSCs into the body, such as under the easily accessible back skin, will result in MSCs autonomously migrating from the scaffold to the site where cytokines are produced, and enhance tissue repair [7]. Collagen gel, which has high biocompatibility and is easily applied in the clinic, was used as a scaffold to administer a sufficient number of stem cells in this study. We investigated the migration of delivered stem cells and healing of a tooth extraction site, where both soft and hard tissue healing are involved.

For current cell therapies, MSC administration routes include intra-arterial/intravenous, intraperitoneal, and direct administration into tissues and organs. These methods can roughly be classified into two groups: systemic administration and local administration. Systemic administration is a well-documented method [8,9], whereby MSCs are applied in the cardiovascular system and reach the damaged area through circulating blood. Cells release various cytokines in a paracrine manner, some of which promote healing of the damaged tissue [10–12]. However, reportedly less than 10% of administered MSCs accumulate in the site of damage, with many cells becoming captured in the lungs [13–15].

Local administration of MSCs to the injury site has advantages, such as a rapid and localized reaction [16]. However, it involves risks such as cells undergoing apoptosis when administered at high density, or bleeding and secondary damage caused by the administration itself. For these reasons, treatment outcomes are not always stable and the ideal administration method has not yet been established for MSC-based therapies.

## Materials and Methods

### 1. Animals

Male C57BL/6 N mice (6 weeks of age; Kyudo Laboratories, Tosu, Japan) and male green fluorescent protein (GFP)-transgenic C57BL/6N mice (CAG-EGFP; 6 weeks of age; Japan SLC, Shizuoka, Japan) were used in this study. All animal experiments were performed under an institutionally approved protocol for the use of animals in research at Kyushu University (approval number: A29-237-0).

### 2. Isolation and culture of MSCs

MSCs were isolated from the bone marrow of mice as described previously. Briefly, bone marrow cells were flushed out from the femoral and tibial bone cavities of the mice. The cells were passed through a 40-µm cell strainer to obtain a single-cell suspension, which was seeded in 100-mm culture dishes at 1 × 10^6^ cells/dish. One day after seeding, cells were washed with phosphate-buffered saline (PBS) and cultured in growth medium comprising Alpha-Minimum Essential Medium (A-MEM; Invitrogen, Grand Island, NY, USA) supplemented with 20% fetal bovine serum (FBS; Equitech-Bio, Kerrville, TX, USA), 2 mM L-glutamine (Invitrogen), 100 U/mL penicillin (Invitrogen), and 100 µg/mL streptomycin (Invitrogen).

### 3. Osteogenic differentiation assay

To confirm calcium deposition, MSCs were passaged on 35-mm dishes at 2.5 × 10^5^ cells/dish, grown to confluence in growth medium, and then cultured in osteogenic culture medium (growth medium containing 1.8 mM KH_2_PO_4_ and 10 nM dexamethasone; both from Sigma-Aldrich, St. Louis, MO, USA). After 28 days of osteogenic induction, cultures were stained with 1% Alizarin Red S solution (Sigma-Aldrich).

### 4. Adipogenic differentiation assay

MSCs were passaged on 35-mm dishes at 2.5 × 10^5^ cells/dish, grown to confluence in growth medium, and then cultured in adipogenic culture medium (growth medium containing 0.5 mM isobutylmethylxanthine, 60 µM indomethacin, 0.5 µM hydrocortisone, and 10 µg/mL insulin; all from Sigma-Aldrich). After 14 days of adipogenic induction, cultures were stained with Oil Red O. The Oil Red O-positive lipid droplets were observed using an inverted microscope (BZ-9000; Keyence, Osaka, Japan).

### 5. CFU-F assay

The CFU-F assay was performed as described previously. Passage 1 MSCs were seeded into culture dishes (Nalge Nunc, Rochester, NY, USA). After 16 days of culture, cells were stained with a solution of 0.1% toluidine blue and 2% paraformaldehyde. Total colony numbers were counted per dish. Three independent experiments were performed.

### 6. MSC injection via the tail vein

GFP-MSCs isolated from the bone marrow of transgenic mice were cultured and passaged three times before injection. Mice (n = 5 per group) were anesthetized by combined anesthesia (0.3 mg/kg medetomidine, 4.0 mg/kg midazolam, 5.0 mg/kg butorphanol) and *ex vivo*-expanded MSCs (1 × 10^6^ cells in 2 mL of PBS) were administered via the tail vein 48 hours before tooth extraction. Control mice (n = 5) were injected with PBS via the tail vein.

### 7. MSC injection into the back of the mice

GFP-MSCs and mice were prepared as above. The *ex vivo*-expanded MSCs (1 × 10^6^ cells in 2 mL of PBS) were then administered into the back of the mice 48 hours before tooth extraction. Control mice (n = 5) were injected with PBS via the same route.

### 8. Cell cultures

For *in vitro* assessment, MSCs were seeded on 35-mm dishes at 2 × 10^4^ cells/dish and incubated for 12 hours at 37°C under 5% CO_2_. A subset of MSCs were incubated in medium containing 100 nM chloromethyltetramethylrhodamine (MitoTracker Orange CMTMRos; Molecular Probes, Eugene, OR, USA) for 20 minutes at 37°C in the dark. After three washes with medium, these cells were examined under an inverted fluorescence microscope (BZ-9000).

### 9. Collagen gel culture

MSCs were embedded three-dimensionally in collagen gel. Neutralized collagen solution was prepared by mixing eight volumes of Cellmatrix type I-A (Nitta Gelatin, Osaka, Japan) with one volume of 10× A-MEM and reconstitution buffer (2.2 g NaHCO_3_ in 100 ml of 0.05 N NaOH and 200 mM HEPES). The solution contained type I collagen at a final concentration of 2.4 mg/mL. To prevent cell attachment and growth directly on cell culture dishes, 1 mL of plain collagen solution was poured into each culture dish as a base layer. MSCs were suspended in neutralized collagen mixture at a density of 3.0 × 10^6^ cells/mL of gel. After allowing the base layer to solidify with a short incubation at 37°C, 1 mL of neutralized collagen mixture containing cells was added. After the top gel layer was formed by additional incubation at 37°C for 15 minutes, 1.5 mL of culture medium (A-MEM with 2.5% FBS and antibiotics) was added into the culture dish. The medium was changed every 3 days [17,18].

### 10. Counting of GFP-MSCs in collagen

To isolate the GFP-MSCs from 3D culture with collagen gel, the collagen gel was dipped into a digestion solution with 10× α-MEM, dispase II (Godo Shusei, Tokyo, Japan), collagenase L (Nitta Gelatin Inc., Osaka, Japan), and NaHCO_3_ (nacalai tesque, Kyoto, Japan) for 30 minutes at 37℃. The MSCs were then collected by centrifuging using a table top centrifuge (KUBOTA Corporation, Tokyo, Japan) at 367 × *g* for 5 minutes, and counted using a TC20 Automated Cell Counter (Bio-Rad Laboratories, CA, USA).

### 11. Collagen gel with MSC injection

To prepare a homogenous mixture, MSCs (1.0 × 10^6^ cells) were stirred on ice with a mixed solution of 1.6 mL of collagen gel (Cellmatrix type IA) and 0.4 mL of reconstitution buffer. Next, mice (n = 5 per group) were anesthetized using a combination anesthetic (described above), and 2 mL of collagen gel containing MSCs was injected into the back of the mouse using a 5-mL syringe (Terumo Corporation, Tokyo, Japan) and 25-G injection needle (Terumo). MSCs were administered at the intersection between the straight line connecting the median line and the shoulder blades on both sides of the mice.

### 12. Counting of injected MSCs in collagen gel

Mice were euthanized under combined anesthesia and the scaffold was removed from the back of the mice. The excised scaffold was placed in a cell culture dish (60 mm × 15 mm) and cultured in digestion solution with 10× α-MEM, dispase II (Godo Shusei), collagenase L (Nitta Gelatin Inc.), and NaHCO_3_ (nacalai tesque) and allowed to permeate and dissolve at 37°C for 30 minutes. After dissolving the scaffold, the cells were examined under an inverted fluorescence microscope (BZ-9000; Keyence).

### 13. Mouse tooth extraction model

Mouse maxillary first molars were extracted under a combination anesthetic. Three days after tooth extraction, mice were euthanized under combination anesthesia. As controls, mice were subjected to tooth extraction but not administered MSCs. Intact maxilla, kidney, liver, lung, spleen, femur, and tibia were harvested *en bloc*.

### 14. Tissue preparation

For *in vivo* assessment, tissues were prepared according to our previously described methods [19,20]. At the end of each experimental period, mice were euthanized under combination anesthesia and perfused intracardially with heparinized PBS, followed by 4% paraformaldehyde (pH 7.4). Maxillae were demineralized in 5% EDTA for 4 days at 4°C. Prepared sites were cut into 10-µm buccopalatal sections with a cryostat at −20°C.

### 15. Immunohistochemistry

Immunofluorescence staining of *in vitro* and *in vivo* samples was carried out by blocking samples with normal serum (matched to the secondary antibody) for 1 hour, followed by incubation with mouse anti-rat GFP and CD90 antibodies (1:100; Sigma-Aldrich) overnight at 4°C. Samples were then treated with a fluorescein isothiocyanate-conjugated secondary antibody (1:200; Jackson ImmunoResearch, West Grove, PA, USA) for 1 hour at room temperature and mounted with 4′,6-diamidino-2-phenylindole (DAPI). A subset of sections were stained with Azan stain [21].

### 16. Statistical analysis

Data are expressed as mean ± standard deviation (SD). Student t-test (Fig. 3B) or one-way analysis of variance with Fisher’s least-significant difference test was performed. Values of P < 0.05 were considered significant. Experiments were performed using triplicate samples and repeated three or more times to verify their reproducibility.

## Results

### 1. MSC behavior in collagen gel scaffold (*in vitro* study)

Figure 1A shows the experimental schedule and method used in the study. MSCs could remain alive in the scaffold for at least 21 days. However, the number of MSCs remaining in the scaffold at 21 days was significantly decreased compared with other days (Fig. 1B). During the first 14 days, MSCs retained their original characteristics, including abilities for self-renewal and multi-directional differentiation. As shown in Figure 1C, MSCs maintained their expression of MSC markers (CD90 and CD105), could be differentiated into adipocytes or osteoblasts based on Oil Red O or Alizarin Red S staining, respectively, and formed colonies from single cells. These data indicated that the limit of MSC stemness retention may be up to 14 days.

**Fig. 1.**
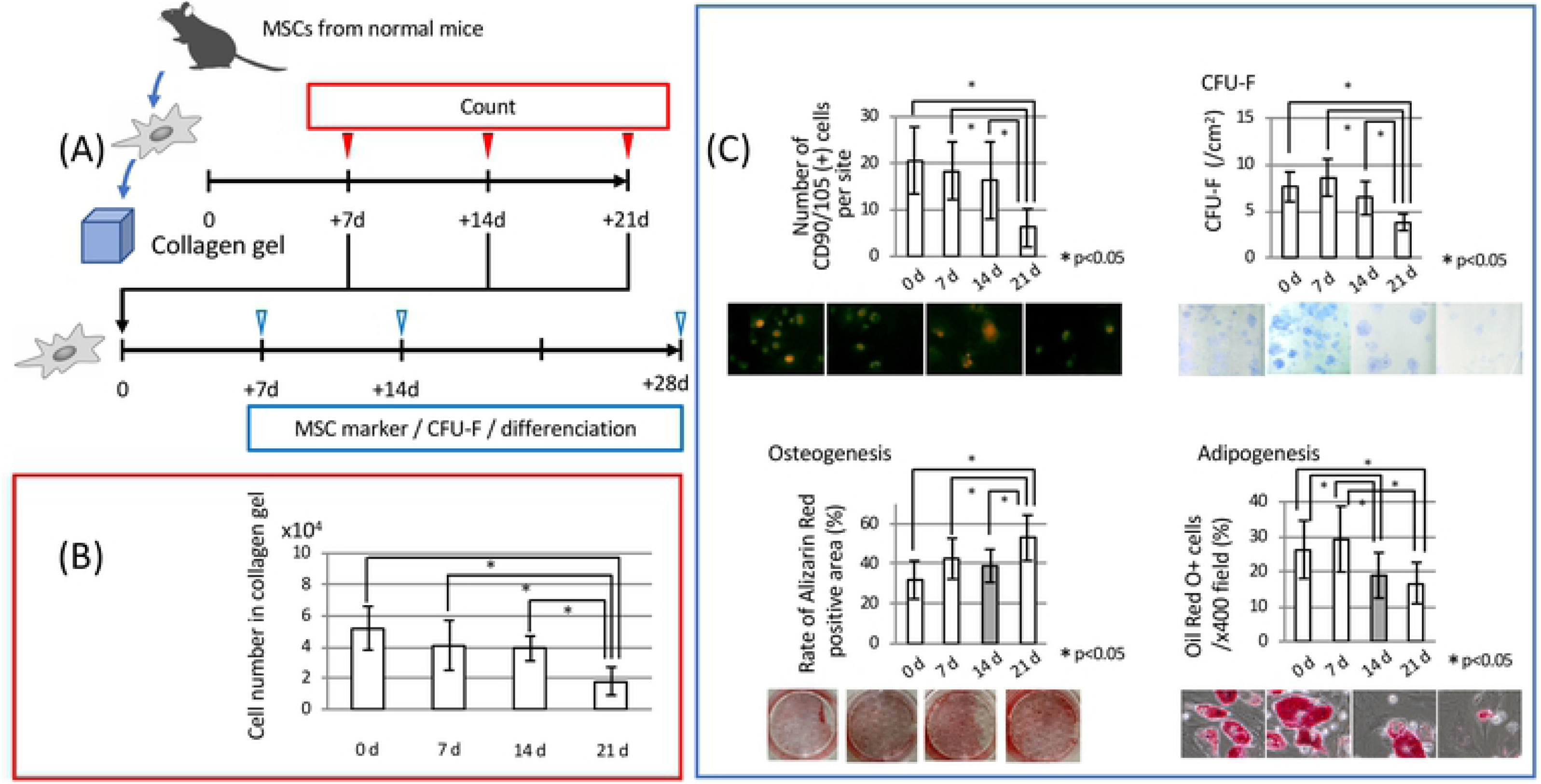
MSC state in collagen gel. **(A)** Experimental protocol for the *in vitro* study. MSCs isolated from wild-type C57BL/6N mice were cultured in a collagen gel for 3 weeks. **(B)** Numbers of surviving MSCs in the scaffold. After the 3-week culture in collagen gel, numbers of cells were evaluated. MSCs could remain alive for at least for 14 days. **(C)** Maintenance of MSC characteristics. After the 3-week culture in collagen gel, isolated cells were checked for the characteristics of MSCs, including self-renewal, multi-directional differentiation, and expression of MSC original markers (CD90 and CD105). (Top left) MSC markers (CD90 and CD105). There were many CD-90/CD-105 double-positive cells for 14 days. (Top right) CFU-F of MSCs. MSCs cultured for 21 days generated fewer CFU-Fs compared with MSCs at other time points. (Bottom left) Osteogenic differentiation of MSCs. MSCs cultured under osteogenic differentiation conditions for 4 weeks were stained with Alizarin Red S. Scale bars: 50 µm. (Bottom right) **Adipogenic differentiation of MSCs**. MSCs cultured under adipogenic differentiation conditions for 2 weeks were stained with Oil Red O. All data indicated that stemness may be retained for up to 14 days. \**p* < 0.05.

### 2. Stability of collagen gel in the body (*in vivo* study)

MSCs were injected with collagen gel into the subcutaneous tissue of mice. The appearances of mouse backs immediately, 7 days, and 14 days after collagen injection are shown in Figure 2B. During this period, the collagen retained its shape with no significant changes. Additionally, as shown in Figure 2B, the gel showed some changes in the number of living cells over 2 weeks, based on GFP/CD90-positive cells (Fig. 2C). Thus, MSCs could be retained in the scaffold for at least 2 weeks *in vivo*, similar to *in vitro* results.

**Fig. 2.**
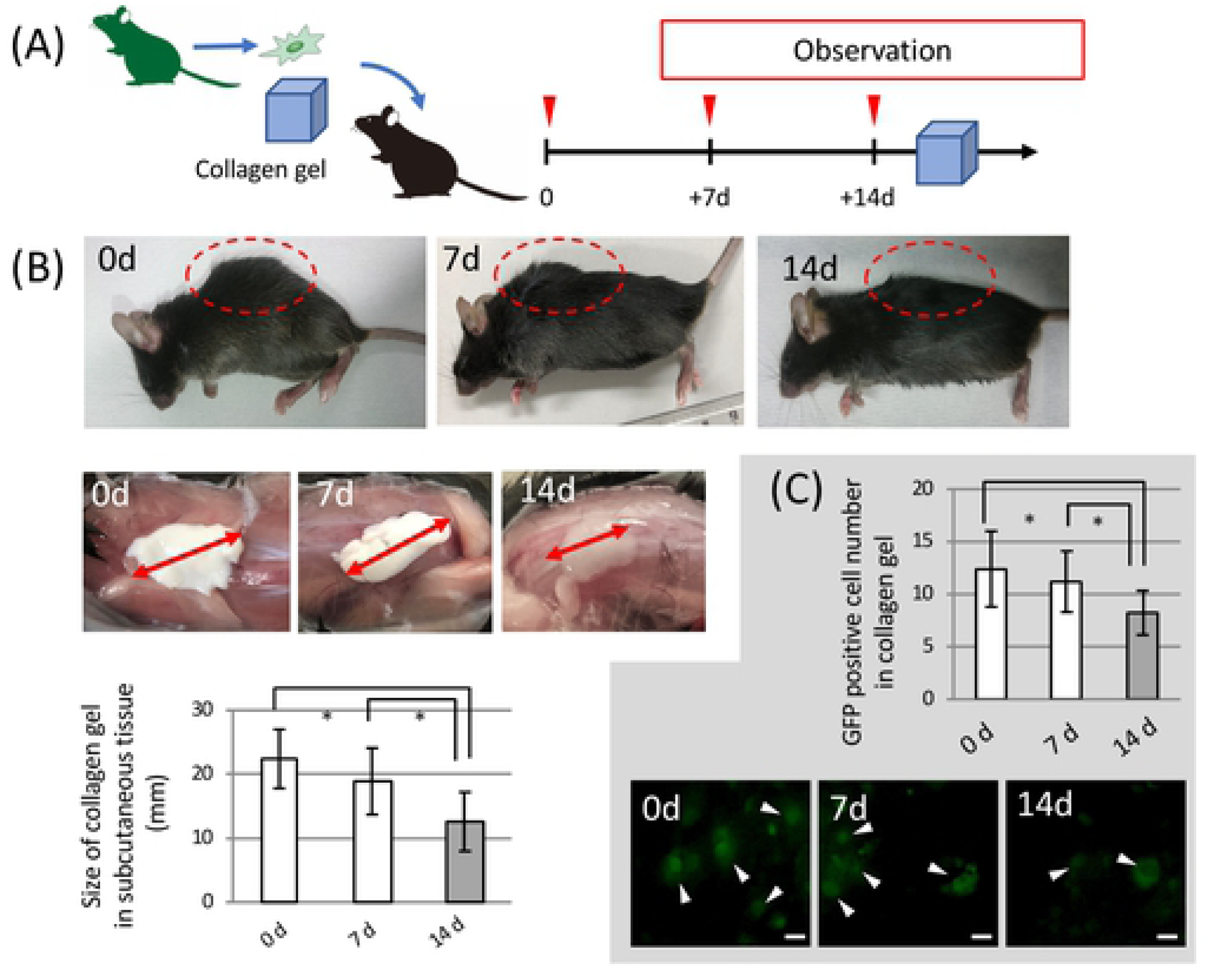
MSC stability in a subcutaneously implanted collagen scaffold. (A) Experimental protocol for *in vivo* experiments. (B) (Upper) Appearances of mouse backs after collagen injection (immediately, 7 days, and 14 days after injection). (Lower) Shapes of collagen gels in the backs of mice (immediately, 7 days, and 14 days after injection). The size of the collagen gel did not change at any time point examined. (C) Numbers of living cells in collagen gels at time of injection and upon removal from mouse backs at 2 weeks after injection. More than half of cells remained alive for 2 weeks. Scale bar, 50 μm. \**p* < 0.05.

### 3. Changes in cell number in scaffolds before or after tooth extraction

As shown in Figure 2, MSCs were retained in the scaffold for about 2 weeks. However, after tooth extraction, the number of MSCs in the scaffold showed a significant decrease compared with the number before extraction (Fig. 3). In particular, at 5 and 7 days after collagen injection with MSCs, there was a large difference between extraction and non-extraction groups.

**Fig. 3.**
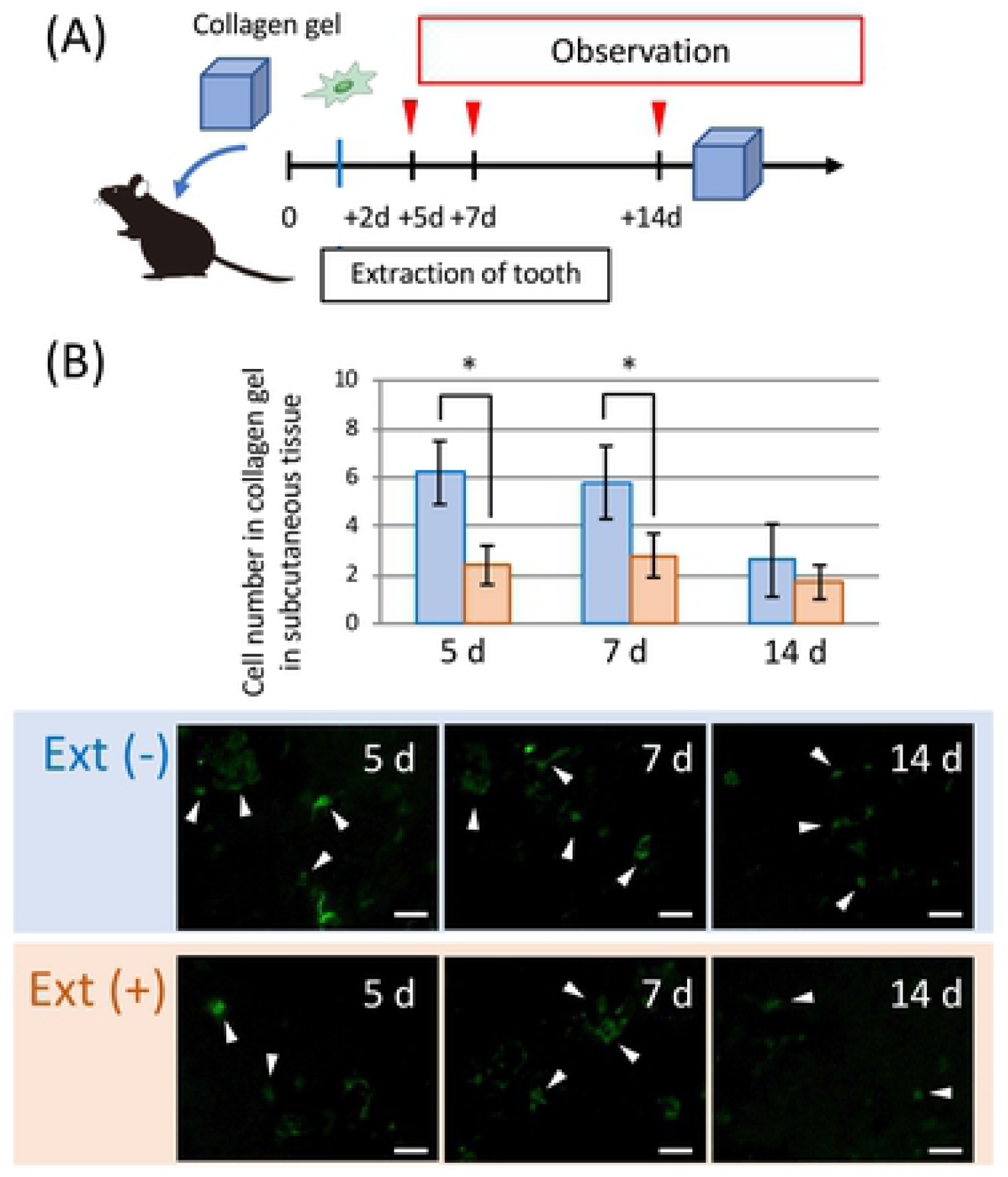
Changes in cell number in the scaffold after tooth extraction. Wild-type C57BL/6N mice received tooth extraction 2 days after collagen injection with MSCs. After a further 3, 5, or 12 days, numbers of cells in the collagen gel were compared between extraction and no-extraction groups. In the extraction group (5 and 7 days), a remarkable reduction of cells in the scaffold was observed. Scale bar, 50 μm. \**p* < 0.05.

### 4. Accumulation in several organs after MSC administration

Numbers of accumulated MSCs in lung, kidney, liver, spleen, and bone marrow after MSC injection via the tail vein are shown in Figure 4B. In particular, the lung accumulated large numbers of MSCs through the blood capillaries. As shown in Figure 4C, we compared differences between tail vein and scaffold injections by countin numbers of accumulated MSCs in the lung. Mice injected with scaffold had much lower numbers of MSCs in the lung than mice injected with MSCs via the tail vein or via the back.

**Fig. 4.**
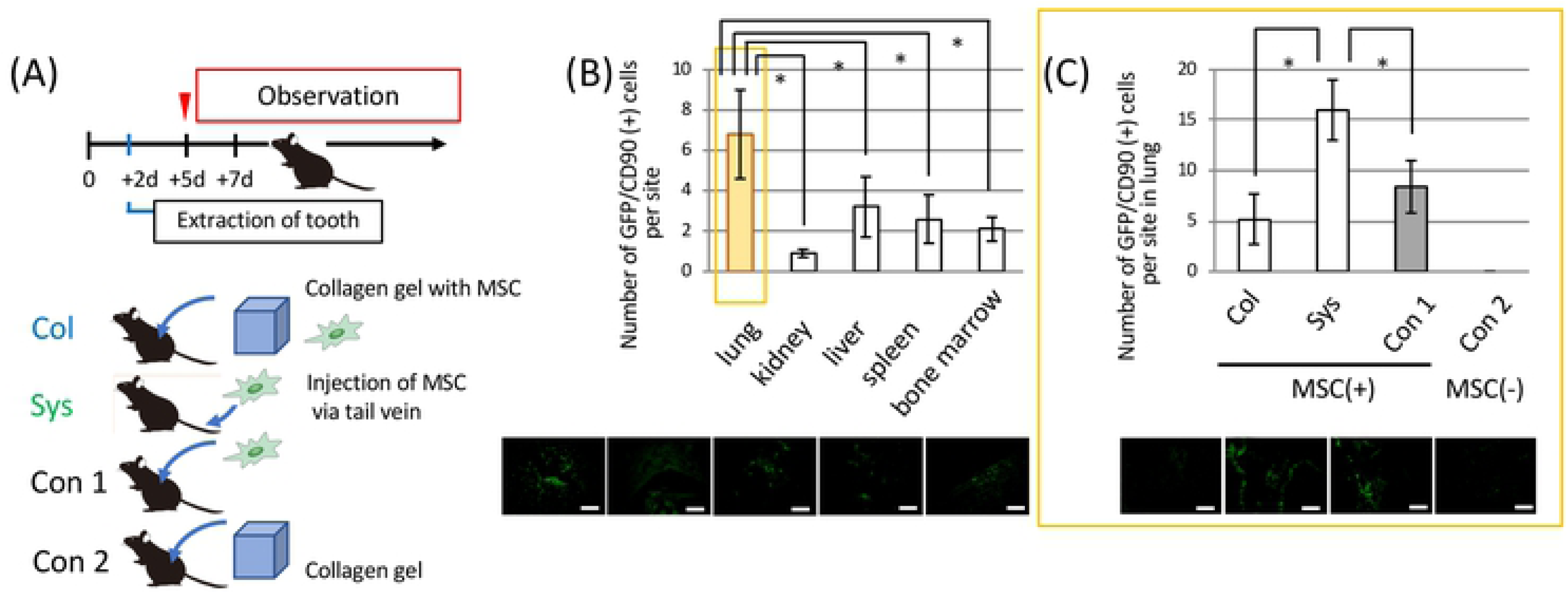
Accumulation of MSCs into organs following tooth extraction. (A) Experimental protocol for *in vivo* experiments. (B) Numbers of MSCs in several organs. At 5 days after systemic injection via the tail vein, cell numbers in the lung, heart, kidney, liver, and spleen were counted and plotted. In particular, the lung accumulated a large number of MSCs. Scale bar, 100 μm (C) As novel and conventional methods, MSCs isolated from GFP-transgenic mice were cultured with or without a collagen gel, and injected into mice 2 days before tooth extraction. At 5 days after MSC administration, numbers of cells accumulated in the lung were evaluated. Systemic injection via the tail vein and injection of MSCs into the back of mice resulted in significantly higher numbers of MSCs in the lung compared with the collagen group. Scale bar, 100 μm. \**p* < 0.05.

### 5. Effect of delivering MSCs in a scaffold on epithelial healing after tooth extraction

In mice injected MSC via the tail vein and in the scaffold MSC groups, many GFP/CD90 double-positive cells were observed around the socket after tooth extraction (Fig. 5B). The numbers of MSCs were similar in the scaffold MSC and tail vein-injected MSC groups. Nevertheless, while the control group (no treatment, injection of MSCs into the back of the mice) exhibited poor healing, the scaffold MSC group showed good mucosal healing, similar to the injected MSC group (Fig. 5C). Furthermore, immunohistochemical staining of Ln-332 revealed the degree of epithelial healing (Fig. 5D). To observe the primary epithelial closure of the tooth extraction cavity, Ln-332 was stained and the continuity of the epithelium was observed. The data indicated that MSC adaptation within a scaffold elicited similar healing ability to MSC adaptation after injection.

**Fig. 5.**
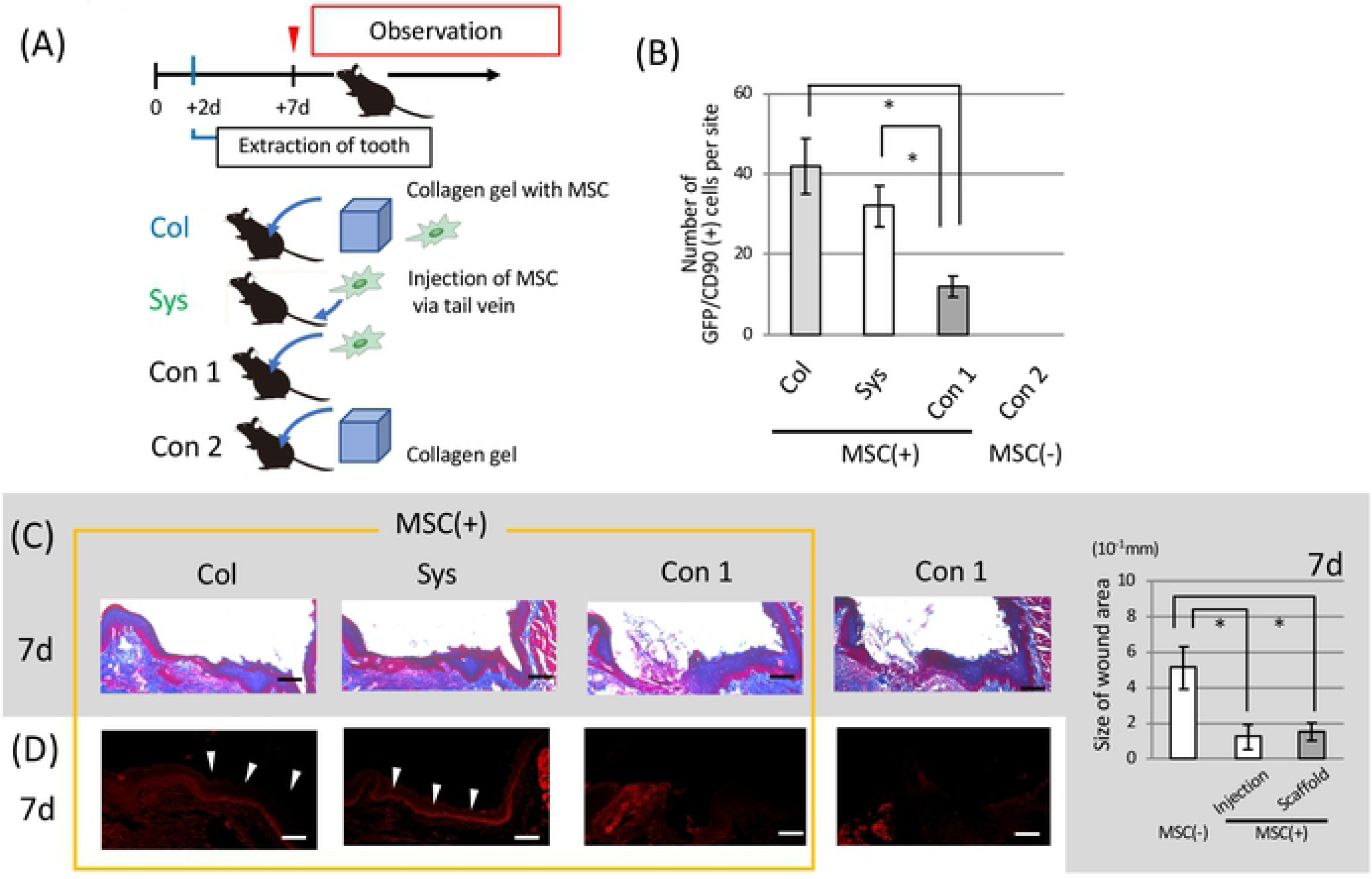
Effect of MSCs from collagen gel for tooth extraction. (A) Experimental protocol for *in vivo* experiments. (B) At 7 days after MSC administration, numbers of accumulated MSCs around the tooth extraction socket. MSCs injected via the tail vein and MSCs from collagen gel were observed as many GFP/CD90 double-positive cells around the socket after tooth extraction by immunofluorescence staining. Injected MSCs accumulated similarly to scaffold MSCs. (C) Wound healing period. The duration of healing was remarkably shortened by both systemic injection and administration in a collagen scaffold. Scale bars: 100 µm. The data indicated that MSC adaptation with a scaffold had similar healing ability to MSC adaptation after injection. (D) Expression of adhesion protein. Laminin-332 was observed as a line on connective tissue in both systemic injection and collagen scaffold groups (white arrow heads). Scale bars: 100 µm. \**p* < 0.05.

## Discussion

Recently, clinical research of stem cell therapies has been increasing. Although principle mechanisms underlying the therapeutic benefit of stem cells remain unclear, many reports suggest the efficacy of related cell therapies [22,23]. In this study, two mechanisms of enhanced healing of the tooth extraction site can be assumed. First, administered MSCs differentiate into epithelial cells and facilitate wound closure. Second, MSCs secrete cytokines and growth factors that activate epithelial cells to migrate and proliferate to achieve primary closure of the damaged tissue.

We previously reported that intravenous administration of MSCs in rats contributes to the healing of damaged tissue [24]. However, delivering cells under the back skin without a scaffold did not result in a positive effect on tissue healing [25]. As such, we focused on the scaffold as a foundation for cells.

First, we searched for a suitable scaffold material in which MSCs could be retained for long periods of time. Although we verified the competence of other materials (data not shown), we selected collagen as a scaffold because it is solid at low temperatures, which makes it easy to deliver by injection and less damaging to cells. In addition, collagen gelates at body temperature, which facilitates retention at the application site [26]. Generally, if artificial materials exist in the body for a long time, problems such as infection or degradation may arise; thus, the characteristics of type I collagen, which can be absorbed by the body, might be suitable. In addition, collagen seems to provide an effective scaffold with little toxicity, as suggested by other studies [27,28]. As shown in Figure 1, MSCs were cultured in the collagen scaffold for 2 weeks, at which time we confirmed their survival and retention of “stemness”.

Consistent with previous reports [29,30], MSCs could remain alive in the scaffold without major changes in cell number for 2 weeks (Fig. 1B). Meanwhile, as shown in Figure 1C, MSCs maintained their expression of MSC markers (CD90 and CD105), and MSCs extracted at each stage exhibited differentiation and self-renewal potential. Therefore, MSCs could maintain their “stemness”, an important characteristic, for at least 14 days.

Second, we checked the competence of collagen as a scaffold. As shown in Figure 2A, the injected 2 mL of collagen retained its form at 2 weeks after insertion into back subcutaneous tissue. Our results were consistent with previous reports suggesting that a 2-week period was required to absorb approximately 5-mm diameter (corresponding to 2 mL) of collagen containing MSCs after insertion into the body as a scaffold for exogenous cells [31,32]. Safety of the collagen scaffold was determined by the survival rate of MSCs in the scaffold in *in vivo* experiments (Fig. 2B). As a result, MSCs in collagen gels implanted in back subcutaneous tissue remained alive for at least 2 weeks, similar to the *in vitro* data shown in Figure 1. Thus, the injected collagen scaffold had the potential to work as a base for exogenous MSCs in the body.

Third, we investigated whether MSCs in the collagen gel could maintain their chemotactic properties. A previous report indicated that systemically delivered MSCs accumulated at sites of inflammation [33]. As shown in Figure 3, MSCs could be guided from the collagen scaffold to the wound area after tooth extraction. However, most MSCs remained in the scaffold when tooth extraction was not performed. MSCs in the collagen gel seemed to have the ability to accumulate via systemic circulation. Therefore, we examined the localization of MSCs after they migrated from the collagen scaffold, as shown in Figure 4.

Regarding the effect of administration method, we previously conducted a study comparing systemic and topical administration, which suggested that subcutaneous or intramuscular administration of MSCs was ineffective when not applied with a scaffold [25]. Similar to previous findings [34,35], the lung and liver had accumulated many cells from the systemic circulation. In this experiment, many MSCs were observed in the lung following systemic and local administration without using scaffolds, but there was only slight accumulation in the lung when the MSCs were injected within a collagen scaffold (Fig. 4C). After tooth extraction, MSCs from the collagen scaffold were not observed in the lung, but could be seen in the wound area. Therefore, delivering MSCs within a scaffold may decrease the risk of lung or cardiac infarction. As MSCs are originally adherent cells, they readily form cell aggregates immediately after isolation from culture dishes [36]. Thus, when injected via the tail vein, most of the cells formed a mass. This might be one reason that intravascularly applied cells have higher risks for capture in capillary-like lung tissue. In contrast, cells migrating from the scaffold should move individually, making them less likely get clogged in the lung, which is supported by our results.

Finally, we investigated the effect of MSCs with a collagen scaffold. As shown in Figure 5B, we examined whether MSCs could accumulate in the tissue around the tooth extraction socket. As mentioned before, many reports suggest that MSCs could accumulate into the wound area after systemic administration [37]. However, since the number of accumulated cells was generally very large in systemic cases, the cells could easily undergo apoptosis around the area [38]. Indeed, as shown in Figure 5B, many cells were observed around the extraction site by conventional systemic injection, but a similar number of cells were observed around the site in the scaffold group. Both groups showed similar promotion of wound healing after tooth extraction, compared with the control group (no treatment, injection of MSCs into the back of the mice) (Fig. 5C).

The results of this study suggested that systemically administered MSCs accumulated and stayed in the lungs, and then moved from there to the site of inflammation to promote tissue healing. In contrast, the high density of MSCs at the injected site following local administration meant that it was hard for them to move from the administration site to the site of inflammation. In addition, the number of MSCs accumulated in the extraction socket was lower because the cells differentiated and their ability to migrate to the inflammation site decreased. As a result, tissue healing was significantly lower following local, compared with systemic administration. In the scaffold group however, the presence of the cells in a suitable space in the collagen apparently maintained their MSC properties. This indicates that side effects such as pulmonary embolism may be avoided by using collagen scaffolds, and that this process may also contribute to tissue healing.

Amounts of cytokines released are reported to closely correlate with the degree of inflammation [39]. In addition, cell migration in response to cytokines has been shown to occur in a dose-dependent manner [40]. This suggests that when severe damage occurs in a tissue, increased inflammation and cytokine release elicit migration of MSCs to that area. Based on this concept, our delivery method should control the number of MSCs migrating to the damaged site depending on the degree of damage, which may overcome some of the disadvantages of existing cell delivery methods.

## Conclusions

After receiving a signal for assistance from the injured site, MSCs transplanted in a scaffold can decide on the number and timing of cells required at the site of injury. This novel method has huge potential for both the treatment and prevention of tissue damage, given the ability of MSCs to detect abnormal and damaged cells. MSC-based therapy using a collagen scaffold may thus offer a safe and effective therapeutic modality for various systemic diseases.

## Acknowledgments

Not applicable

## Author Disclosure and Ghostwriting

All authors declare no competing financial interests exist. The authors listed expressly wrote the content of this article. No ghostwriters were used to write this article.

## About the authors

NU, IA, YA, IN, and YM were involved in the practical aspects of experiments. NU, IA, YA, TY, RK, YW and XZ collected, analyzed, and interpreted the data. IA designed the study and provided financial and administrative support. IA, YA, TY, AF and KK wrote the manuscript. IA, TY, AF and KK revised the manuscript for publication. Each author participated sufficiently in the work to take public responsibility for appropriate portions of the content.

## Abbreviation

A-MEM: Alpha Minimum Essential Medium
BSA: bovine serum albumin
CFU-F: colony forming unit-fibroblast
DAPI: 4’,6-diamidino-2-phenylindole
Dex: dexamethasone
FBS: fetal bovine serum
FITC: fluorescein isothiocyanate
GFP: green fluorescent protein
MSC: mesenchymal stem cell
PBS: phosphate-buffered saline

